# Electronic Cigarette Aerosol Exposure Induces Airway Remodeling in 3D Human Tracheobronchial Epithelial Tissues: From Goblet Cell Hyperplasia to Squamous Metaplasia

**DOI:** 10.1101/2025.10.30.685605

**Authors:** Rattapol Phandthong, Edgar Rolon, Brianna Ramirez, Mohamad Kudsi, Audrey Choi, Arash Tamasbi, Teresa Martinez, Prue Talbot

**Affiliations:** Department of Molecular, Cell and System Biology, University of California, Riverside, CA 92521, USA

**Keywords:** Electronic Cigarette, Goblet Cell Hyperplasia, Squamous Metaplasia, Airway Remodeling Involucrin, Cilia, 3D Human Tracheobronchial Epithelial Tissue

## Abstract

Electronic cigarette (EC) aerosols are increasingly recognized for their disruptive effects on airway physiology, yet the temporal progression of epithelial remodeling remains poorly characterized. While squamous metaplasia, mucin hypersecretion, and ciliary dysfunction are hallmark features of smoking-related airway disease, analogous data for EC exposure are limited. To address this gap, we employed a 3D human tracheobronchial epithelial tissue (hTET) model derived from airway basal stem cells and exposed it daily to NJOY ACE “Classic Tobacco” aerosols (1 or 5 puffs) or clean air for 1, 2, or 3 weeks. Remodeling was assessed via immunolabeling and Western blot analysis. MUC5AC expression revealed a transient hypersecretory response, peaking at week 2 and declining by week 3. Multiparametric analysis demonstrated progressive squamous remodeling: early ciliary loss and involucrin upregulation emerged by week 1, with severity increasing in a puff-dependent manner. By week 2, tissues exhibited marked ciliary depletion, altered epithelial thickness, horizontal nuclear reorientation, and elevated involucrin. At week 3, both exposure groups developed a squamous metaplasia phenotype, characterized by involucrin-positive squamous cells replacing ciliated cells. Quantitative analysis confirmed sustained involucrin elevation, with the 1-puff group reaching levels comparable to the 5-puff group by week 3. These findings delineate a sequential pathological trajectory initiated by EC aerosols, progressing from transient goblet cell activation to squamous metaplasia. This human-relevant model underscores the potential for repeated low-level EC exposure to induce airway epithelial remodeling and pathology.

## Introduction

Since their introduction in the early 2000s, electronic cigarettes (ECs) have rapidly gained global popularity as alternatives to conventional tobacco cigarettes. Marketed as safer options and promoted as smoking cessation aids, ECs have been widely adopted, particularly among young adults [1–2]. However, accumulating clinical, infodemiological, and toxicological evidence has raised substantial concerns regarding their safety [3–8]. Early infodemiological reports, followed by a systematic review of case reports, showed that EC use was linked to a broad range of adverse outcomes across respiratory, cardiovascular, gastrointestinal, neurological, and immune systems, suggesting potential risks even for healthy individuals [7,9]. Building on these early signs of harm, subsequent case reports have described respiratory disorders , including acute lung injury, constrictive bronchiolitis, and lipoid pneumonia [9–13].

Interpretation of EC-related health risks is complicated by the extensive variability in device design, brand-specific formulations, and user topography [14–20]. EC liquids and aerosols contain diverse chemicals, including nicotine, flavor chemicals, solvents, reaction products, and metals, many of which lack comprehensive inhalation toxicology data [21–31]. This chemical heterogeneity presents significant challenges for risk assessment and regulatory oversight.

To date, most health-related studies on ECs have focused on mode of action endpoints, such as cytotoxicity, inflammatory responses, oxidative stress, and perturbations in cellular signaling pathways [32–36]. While these studies provide valuable insight into short-term injury, they are limited in their ability to define the specific diseases associated with long-term EC use. Unlike conventional cigarette smoking, which has been conclusively linked to chronic airway diseases, the long-term disease consequences of EC exposure remain poorly characterized. This knowledge gap is attributable in part to the relatively recent introduction of ECs and the long latency periods required for many progressive diseases to manifest.

Recent findings have begun to address this gap. A meta-analysis study by Glantz et al. [37] reported that exclusive EC use is associated with significantly increased odds of developing some respiratory and metabolic diseases compared to non-users, challenging the perception that ECs are a safe alternative to conventional cigarettes. A complementary longitudinal cohort study found that exclusive use of ECs was associated with new cases of chronic obstructive pulmonary disease (COPD) and, among adults aged 30–70, with hypertension, indicating cardiopulmonary risks related to vaping [3]. In parallel, transcriptomic analyses of nasal airway epithelia from EC users revealed elevated expression of involucrin, keratin 10, and keratin 13, established markers of squamous metaplasia (SM) [38]. However, no prior in vitro human airway model has directly characterized the progressive onset of SM in response to repeated EC aerosol exposure, leaving a critical gap in understanding its initiation and trajectory.

To address this need, we developed a “disease-in-a-dish model” for SM using controlled air liquid interface (ALI) exposure and quantitative assays to monitor its progression in 3D human tracheobronchial epithelial tissue (hTET) produced in vitro from airway basal stem cells. This model enables temporal observation of epithelial remodeling, mucociliary dysfunction, and squamous induction within a 3-week period, providing a tractable platform to study airway disease progression. SM was selected as the condition of interest due to its relevance as a precursor to debilitating airway diseases, including squamous cell carcinoma [39] and its frequent occurrence in individuals with COPD [40,41]. To evaluate SM, hTETs were exposed daily for 3 weeks to aerosols generated from NJOY ACE EC (classic tobacco flavor). Tissues were assessed at 1-, 2- , and 3-week using endpoints that included MUC5AC expression, ciliary loss, ciliary height, increased involucrin expression, epithelial thickness, and presence of horizontally oriented nuclei, all hallmarks of SM progression. These data establish a robust in vitro model for investigating pathological changes in the respiratory epithelium during exposure to environmental chemicals, including EC aerosols. The model offers utility for evaluating other EC formulations, flavoring agents, and chemical constituents, and may serve as a platform for regulatory toxicology and disease-relevant screening.

## Materials and Methods

### Human Airway Basal Stem Cell (ABSC) Isolation

A de-identified human lung from a female donor was obtained from the International Institute for the Advancement of Medicine (IIAM.org). Airway basal stem cells from the tracheobronchial region were isolated using established procedures [42–44]. Isolated cells were seeded onto collagen type I–coated T-75 flasks at 5,000 cells/cm² and cultured in Bronchial Epithelial Cell Growth Medium (Lonza Walkersville CC3170; Cat. No. NC9202780) supplemented with antibiotic–antimycotic solution (Thermo Fisher, Tustin, CA, USA; Cat. No.15240062). ABSCs were maintained at 37 °C, 5% CO₂, and 95% relative humidity. Upon reaching ∼80% confluence, cells were passaged and cryopreserved in liquid nitrogen (−196 °C) until use.

### Human Airway Basal Stem Cell Culture and Differentiation at the Air-liquid Interface

Human ABSCs were cultured in T-25 cell flasks precoated with collagen type 1 (Sigma-Aldrich, St. Louis, MO, USA; Cat. # C89819) and cultured in PneumaCult ExPlus (STEMCELL Technologies, Vancouver, CA; Cat 05040). Cells were seeded at 5,000 cells/cm^2^. At ∼60% confluency, cells were passaged and seeded at 300,000 cell/cm^2^ on the apical side of 12-well transwells cell culture plate (Corning, Inc., Corning, NY, USA; Cat# 3460) precoated with collagen type 4 (Sigma-Aldrich, St. Louis, MO, CA; Cat C7521). PneumaCult ExPlus culture medium at the apical and basal-lateral compartment was changed every other day, until a uniform monolayer formed (>95% density), after which the apical medium was removed, exposing tissue to air, and the medium at the basal-lateral side was replaced with fresh PneumaCult ALI (STEMCELL Technologies, Vancouver, CA; Cat 05001). Cells were cultured at the ALI for 4 weeks to form fully differentiated hTET. PneumaCult ALI medium in the basal lateral compartment was changed every other day. Cells and tissues were cultured in a 37°C, 5% CO_2_, and 95% relative humidity incubator.

### Air-liquid interface Exposure with NJOY Ace “Tobacco Flavor” EC

NJOY Ace EC devices with “Classic Tobacco” pods were purchased online, at local gas stations, and at grocery stores. Aerosol exposures were performed using a Cultex® RFS compact exposure module (Cultex Laboratories GmbH, Hannover, Germany) to deliver to tissue cultures either humidified, sterile air (clean air control), or EC aerosols. For each exposure, tissues were placed in the Cultex exposure chamber that contained culture medium and was maintained at 37 °C. The Cultex® system consisted of a sampling module connected to a custom-designed EC smoking robot (RTI International, NC, USA) and an aerosol guiding module [44,45]. The smoking robot drew either filtered clean air or EC aerosol into a 200 mL syringe inside a biosafety cabinet. Each puff consisted of 55 mL of either filtered air or aerosol, drawn over 4 s, with a 30 s interval between puffs. The collected sample was transferred into the aerosol guiding module, where it was mixed with humidified zero air at 1 L/min to dilute the aerosol and create a uniform flow. This mixture was then directed over the tissue surface, allowed to settle for 5 s, then vented from the chamber at 5 mL/min using a mass flow controller (Boekhorst, Bethlehem, PA, USA) before being discharged into a waste container. hTETs were exposed daily to 1 or 5 puffs of clean air or EC aerosols for 1, 2, or 3 weeks. Between each exposure, tissues were returned to the incubator. After the final exposure, tissues were incubated for 24 h before downstream analyses.

### Tissue Processing

After 1, 2, or 3 weeks of treatment, membranes with hTETs were cut out from the transwells, then washed three times with phosphate-buffered saline containing calcium and magnesium (PBS+; Lonza, Walkersville, MD, USA; Cat. # 17-513F) to remove cell debris and mucous. Tissue was cut into three pieces, then subjected to Western Blotting, <u>b</u>ird’s <u>e</u>ye <u>v</u>iew (BEV) immunofluorescence microscopy, or histology. Samples prepared for Western blotting were lysed with RIPA buffer with phenylmethylsulfonyl fluoride protease inhibitors (ChemCruz Biochemical, Dallas, TX; Cat. No.sc-24948A) then centrifuged at 3,000 rpm to separate cell debris from the protein supernatant. Protein was quantified in the lysate using the Pierce bicinchoninic acid (BCA) assay kit (Thermo Scientific, Tustin CA; Cat. No. 23225). hTETs subjected to immunofluorescence and histology were washed with 10 mM dithiothreitol (DTT with PBS without calcium and magnesium) for 5 minutes to remove additional mucus at the apical surface, then fixed with 4% paraformaldehyde for 30 minutes, followed by three washes with PBS+.

### Bird’s Eye View (BEV) Immunofluorescence Microscopy

Fixed hTETs were permeabilized using 0.01% Triton X (PBS+ with Triton X; Sigma-Aldrich, St. Louis, MO, USA; Cat. # X100) for 30 minutes and incubated in a blocking solution (donkey serum + 0.01% Triton X) for 1 hour. Tissues were incubated in specific primary antibodies diluted in PBS+ 0.01% Tween (PBST; Sigma-Aldrich, St. Louis, MO, USA; Cat. # P1379) overnight at 4°C, then washed three times with PBST. A secondary antibody conjugated to a fluorophore was diluted to a working concentration with PBST and incubated with the tissue for 2 hours, followed by three 10-minute washes in PBST. Tissues were placed on glass slides, embedded in Vibrance® antifade mounting medium with DAPI (Vectashield, Newark, CA, USA; Cat. No. H-1800-10) and protected with coverslips. The tissues were imaged using a Nikon Eclipse fluorescence microscope. Primary antibodies were anti-involucrin (1:500; Proteintech, Rosemont, IL, USA; Cat. # 28462-1-AP), anti-acetylated α-tubulin conjugated to Alexa fluor-594 (1:500; Santa Cruz, Dallas, TX, USA; Cat. # sc-23950 AF594), and anti-MUC5AC (1:500; Thermofisher, Tustin CA, USA; Cat. # MA5-12178). Secondary antibodies were donkey anti-mouse and donkey anti-rabbit, which were conjugated to Alexa Flour 488 Alexa or Fluor 594, respectively (1:500; Thermofisher, Tustin CA, USA; Cat. # A21203, A21206).

### Histological Immunofluorescence

Fixed hTETs were processed in a Shandon Excelsior ES Tissue Processor (Thermofisher, Tustin, CA) at the UCR Histology Core. The hTETs were dehydrated with a graded series of ethanol (70-100%), cleared with CitriSolv (Decon Labs Inc., King of Prussia, PA, USA; Cat.#1601), and infiltrated with melted paraffin at 60°C. The paraffinized tissues were embedded in a block of paraffin then sliced with a microtome at 10 µm to produce sections of hTETs, which were captured on positively charged slides for imaging. After hTET tissues were rehydrated deparaffinized with xylene using two 30-minute baths, followed by rehydration with a graded series of ethanol (100%-30%) baths, two distilled water baths, and a PBS+ bath. Then tissue was placed in a sodium citrate buffer (10 mM sodium citrate, 0.05% Tween 20, pH 6.0) at 32.2°C for 20 minutes, followed by a rinse of water and three 5-minute PBS+ baths. Tissues were permeabilized using 0.01% Triton X (Triton X in PBS+) for 10 minutes, followed by incubation in blocking solution (donkey serum + 0.01% Triton X) for 1 hour. Tissues were labeled with antibodies and imaged as described in the above section on BEV.

### Image analysis

BEV and histological images were analyzed using CL-Quant software (DR Vision, Seattle, WA), which extracts signal area and intensity measurements. ImageJ software was used to measure tissue width and cilia height. CellProfiler software [46] was used to segment the nuclei, using automatic detection with manual assistant to correct errors and determine the orientation angle of the nuclei. All analyses were performed on original images with unmodified intensity values. Representative images were uniformly enhanced in Nikon Elements (Nikon Instruments, Melville, NY, USA) using identical brightness and contrast adjustments for visual clarity.

### Western blotting

Following lysate preparation, denaturing buffer (β-mercaptoethanol and Laemmli, 1;10; Bio-Rad, Carlsbad, CA, USA; Cat. No. 1610710 and 1610747) was added to Western blotting lysate at 1:4 ratio. In brief, buffer-lysate mixtures were heated at 95°C for 1 minute, then loaded onto Any kD™ Mini-PROTEAN® TGX Stain-Free™ Protein Gels (Bio-Rad, Carlsbad, CA, USA; Cat. No. 4568124) for electrophoretic separation of proteins at 120 V in running buffer (diluted to 1X in distilled water from 10x/Glycine/SDS Buffer; Bio Rad, San Francisco, CA, USA; Cat. 161-0772). Proteins were then transferred onto a wet PVDF membrane (Bio Rad, San Francisco, CA, USA; Cat.1620174) at 120 V in transfer buffer (tris base 25nM, glycine 192 nM, and 15% methanol). Membranes were cut horizontally at the expected location of the proteins of interest based on their molecular weight (kDa). Blots were probed with anti-involucrin (1:1000; Proteintech, Rosemont, IL, USA; Cat. # 28462-1-AP) and a secondary anti-rabbit conjugated to HRP (1:1000; Cell Signaling Technology, Danvers, MA, USA; Cat. 7074S). Anti-β-actin was used as a loading control (1:1000; Santa Cruz, Dallas, TX, USA; Cat. sc-47778 HRP). Densitometry analysis was done in ImageJ to quantify protein levels.

## Statistical Analysis

Statistical analyses were done with GraphPad Prism 10.6.1 software (GraphPad, San Diego, CA). If data satisfied the assumptions of ANOVA (homogeneity of variances and normal distribution), they were analyzed using a one-way ANOVA. When the assumptions of ANOVA were not met, log(y) transformed data were used for the ANOVA. When significance was found (ANOVA p < 0.05), group means were compared with the corresponding clean air controls or between exposure groups using Tukey’s multiple comparison post hoc test. Data were plotted as mean ± standard deviations using GraphPad Prism 7 software.

## Results

### EC Aerosol Induced Transient Goblet Cell Hyperplasia Prior to Squamatization

Goblet cell hyperplasia and SM were evaluated over 3 weeks of ALI exposure to EC aerosol using fully differentiated hTETs. NJOY ACE “Classic Tobacco” ECs were selected for aerosol generation because they have FDA premarket authorization and represent a substantial market share in the United States [47]. The exposure protocol was designed to approximate vaping patterns representative of some EC users (Figure 1A). After a 4-week differentiation period, hTET were exposed daily to either humidified clean air (control) or NJOY ACE aerosol at 1 or 5 puffs/day for 1, 2, or 3 weeks (Figure 1A). Goblet cell hyperplasia and SM progression were evaluated using morphological and molecular endpoints.

**Figure 1:**
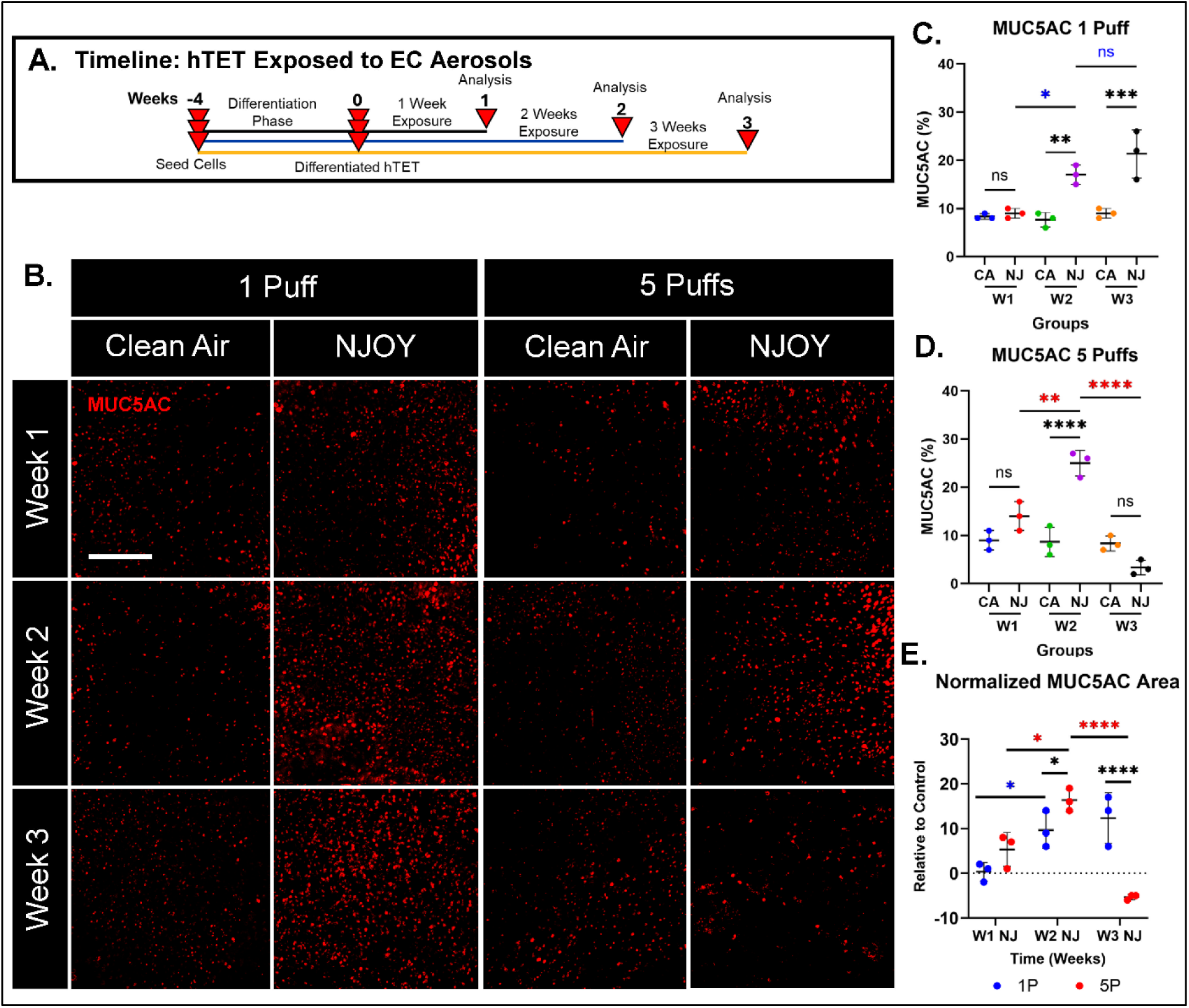
EC Aerosol Induces Transient Goblet Cell Hyperplasia in hTET. **(A)** Experimental timeline showing daily exposure and sample collection after 1, 2, and 3 weeks. **(B)** BEV immunofluorescence micrographs of hTETs exposed daily for 1, 2 and 3 weeks to 1 (left) or 5 puffs (right) of clean air or NJOY EC aerosols. Tissues were labeled with antibodies to MUC5AC. **(C-D)** Graphs showing percentage of MUC5AC area in hTET exposed to 1 puff **(C)** and 5 puffs of EC aerosol **(D)**. **(E)** Normalized MUC5AC area in NJOY-exposed tissues relative to their clean air controls. All graphs show the mean ± standard deviation of three independent experiments. In **C, D,** and **E** log(y) transformed data were analyzed using a one-way ANOVA followed by Tukey’s post hoc test to compare means. Black * or “ns” = comparisons against the clean air control (C, D) and between 1-puff and 5-puff groups (E). Colored * or “ns” = week-to-week comparisons within each exposure group (blue = 1 puff; red = 5 puffs). “ns” = not significant.* = p < 0.05, ** = p < 0.01, *** = p < 0.001. Scale bar = 200µm.

MUC5AC, a well-established marker of goblet cells [48], is frequently elevated in individuals with a history of cigarette smoking [49]. To obtain an overview of goblet cells in hTET exposed to EC aerosol, MUC5AC expression was assessed using immunofluorescence in BEV samples (Figure 1B). In the 1 puff group, fluorescent signal progressively increased over the 3-week exposure period. In contrast, the 5-puff group exhibited a transient increase at week 2, followed by a decline to baseline levels by week 3. Quantification of MUC5AC-positive area as a percentage of total tissue area (Figures 1C and 1D) revealed no significant differences from clean air controls at week 1 in either exposure group. However, by week 2, both groups showed significant increases relative to controls. At week 3, the 1 puff group continued to increase significantly, whereas the 5-puff group returned to control levels.

To enable direct comparisons across exposure groups, all data were normalized to their respective clean air controls (Figure 1E). At week 1, MUC5AC expression did not differ significantly across puff number groups. By week 2, expression was significantly elevated in the 5-puff group relative to the 1-puff group. Conversely, at week 3, the 1-puff group exhibited higher MUC5AC levels than the 5-puff group, which was now similar to the clean air control.

### hTETs Exposed to EC Aerosol for 1 Week had Early Signs of Squamous Metaplasia

The decline in MUC5AC observed in the 5-puff group at week 3 suggested a transition from a hypersecretory phase to an alternative epithelial state. To evaluate whether this shift represented the onset of squamous remodeling, various endpoints were examined after 1 week of exposure (Figure 2). In fluorescent histological sections, there was no significant change in either exposure group in ciliary height (visualized with anti-acetylated tubulin), horizontal nuclei (visualized with DAPI), or tissue thickness (Figure 2 F, G, H), indicating that major structural remodeling had not occurred at this time point. In BEV images, ciliary area showed a modest but significant reduction in the EC 5-puff group, and involucrin integrated density (visualized with anti-involucrin) increased significantly in both EC exposure groups (Figure 2A-C).

**Figure 2:**
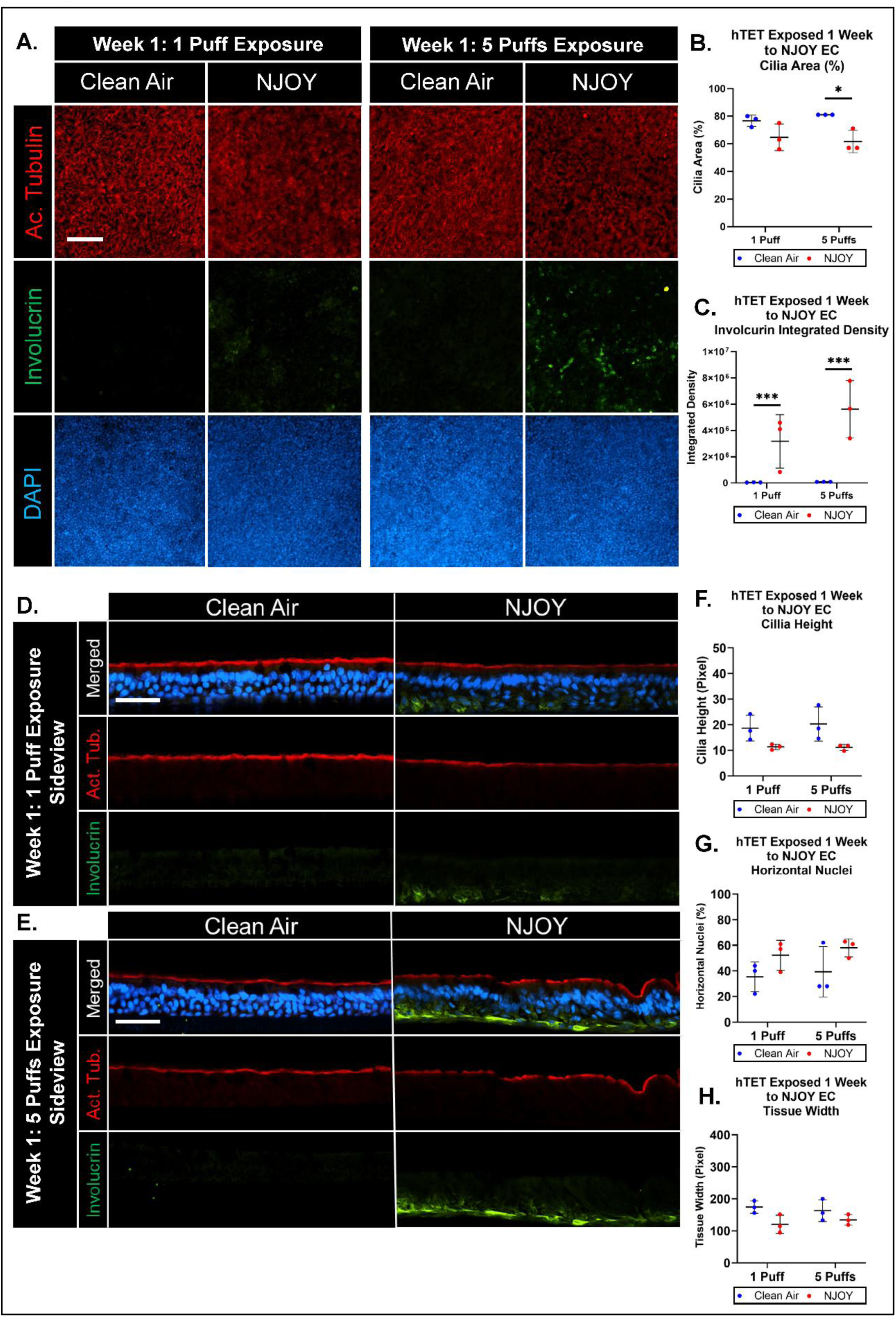
1 Week of NJOY EC Aerosol Exposure Induced an Early Stage of Epithelial Remodeling. **(A)** BEV immunofluorescence micrographs of hTETs exposed daily for 1 week to 1 (left) or 5 puffs (right) of clean air or NJOY EC aerosols. Tissues were labeled with DAPI (nuclei; blue) and with antibodies to acetylated α-tubulin (cilia; red) or involucrin (green). Scale bar = 100 µm. **(B-C)** Graphs showing **(B)** Percent of ciliary area covering the surface of hTETs in BEV images and **(C)** involucrin fluorescence signal as integrated density in BEV images. **(D-E)** Histological sections of hTETs exposed to 1 or 5 puffs of clean air or NJOY EC aerosol. Tissues were stained with DAPI (nuclei; blue) and with antibodies to acetylated α-tubulin (cilia; red) or involucrin (green). Scale bar = 50 µm. **(F-H)** Graphs showing morphometric analysis of **(G)** ciliary height, **(H)** percentage of horizontally oriented nuclei, and **(I)** tissue width. All graphs show the mean ± standard deviation of three independent experiments. In **C, D, G, H** and **I**, log(y) transformed data were analyzed using a one-way ANOVA followed by Tukey’s post hoc test to compare the means. * = p < 0.05

### Two Weeks of EC Aerosol Exposure Produced Additional Evidence of Squamous Metaplasia

Similar assessments were made following 2 weeks of exposure to EC aerosol (Figure 3). In BEV micrographs, tissues exposed to clean air were morphologically normal and intact (Figure 3A). In contrast, tissues exposed to either 1 or 5 puffs of NJOY aerosol had a marked depletion of cilia, evidenced by reduced acetylated tubulin signal in BEV images (Figure 3A). Quantification of BEV data confirmed that ciliary area was significantly reduced in both exposure groups relative to the clean air control with greater loss observed in the 5-puff group compared to the 1-puff group (Figure 3B). Histological analysis further demonstrated significantly decreased ciliary height in both EC-exposed groups (Figure 3F). Involucrin was clearly significantly elevated in BEV images from both NJOY exposure groups relative to clean air controls with the 5-puff group showing significantly higher signal than the 1-puff group, consistent with a concentration-dependent effect (Figure 3A, C). Tissues exposed to 5 puffs displayed elongated cells with squamous morphology and strong involucrin labeling (Figure 3A). Histological micrographs revealed significantly more horizontally oriented nuclei in both EC exposure groups (Figure 3G), along with reduced epithelial thickness, particularly in the 5-puff group (Figure 3H). Taken together, these findings from the 2-week exposure group showed a transition from normal pseudostratified epithelium to tissue with squamous characteristics that included ciliary loss, increased involucrin expression, and altered epithelial architecture.

**Figure 3:**
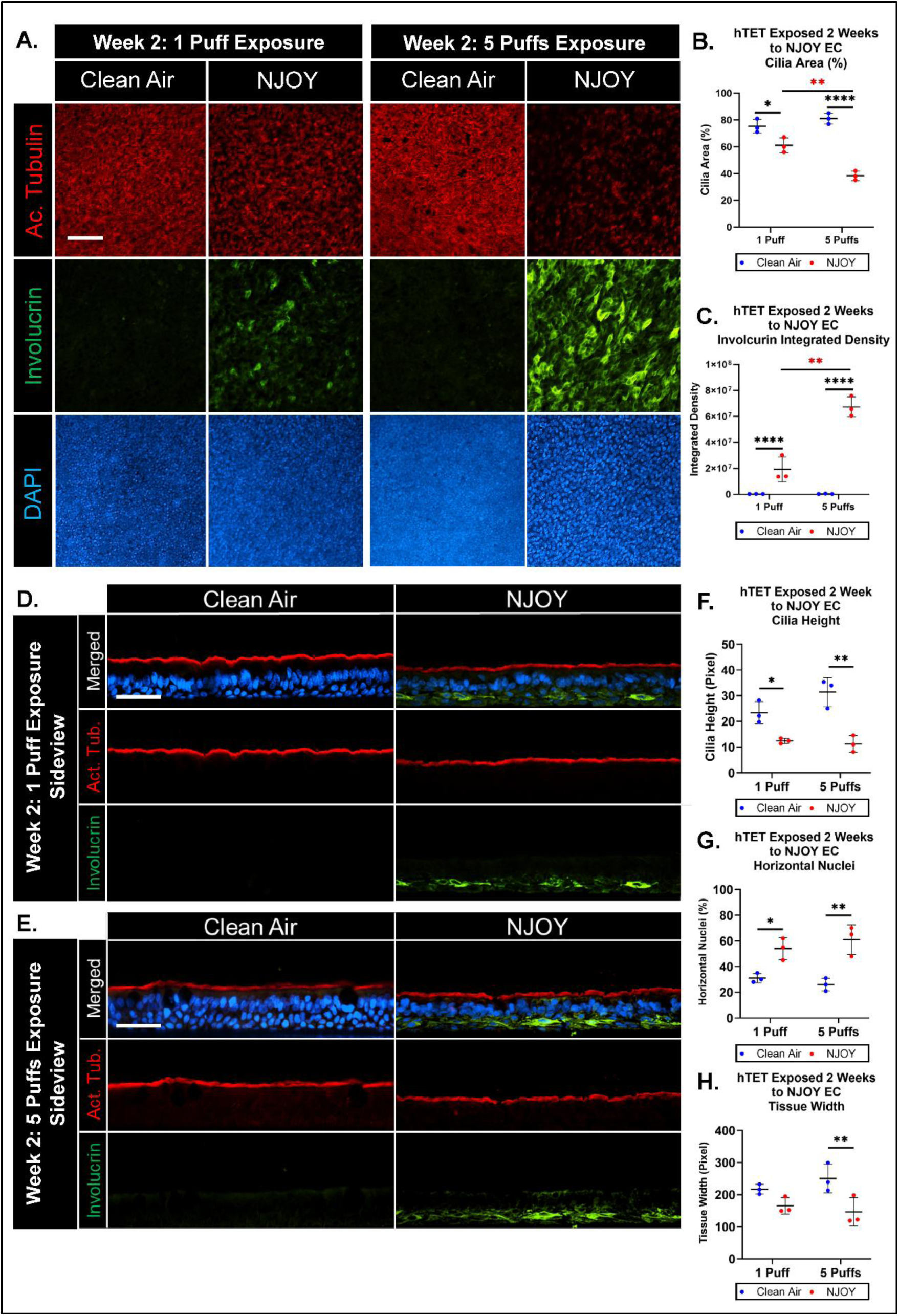
2 Weeks of NJOY EC Aerosol Exposure Induced a Concentration Dependent Transition from Normal Airway Epithelium to Squamous Metaplasia. **(A)** BEV immunofluorescence micrographs of hTETs exposed daily for 2 weeks to 1 (left) or 5 puffs (right) of clean air or NJOY EC aerosols. Tissues were labeled with DAPI (nuclei; blue) and with antibodies to acetylated α-tubulin (cilia; red) or involucrin (green). Scale bar = 100 µm. **(B-C)** Quantification of **(B)** ciliary coverage and **(C)** involucrin fluorescence signal as integrated density. **(D-E)** Histological sections of hTETs exposed to 1 or 5 puffs of clean air or NJOY EC aerosol. Tissues were stained with DAPI (nuclei; blue) and with antibodies to acetylated α-tubulin (cilia; red) or involucrin (green). Scale bar = 50 µm. **(F-H)** Morphometric analysis of histological sections showing **(F)** ciliary height, **(G)** percentage of horizontally oriented nuclei, and **(H)** tissue width. All graphs show the mean ± standard deviation of three independent experiments. In **B, C, F, G,** and **H** log(y) transformed data were analyzed using a one-way ANOVA followed by Tukey’s post hoc test to compare the means. * = p < 0.05, ** = p < 0.01, *** = p < 0.001, **** = p < 0.0001.

### 3 Weeks of EC Aerosol Exposure Induced Squamous Metaplasia

After 3 weeks, hTETs exposed daily to either 1 or 5 puffs of NJOY EC aerosol exhibited pronounced ciliary loss, characterized by diminished acetylated tubulin signal in BEV images and histological sections (Figure 4A, D, E). In contrast, tissues exposed to clean air remained morphologically intact, with well-preserved ciliary architecture. Quantitative analysis confirmed significant reductions in cilia area (BEV images) and ciliary height (histological sections) in both EC aerosol exposure groups relative to controls (Figures 4B, F).

**Figure 4:**
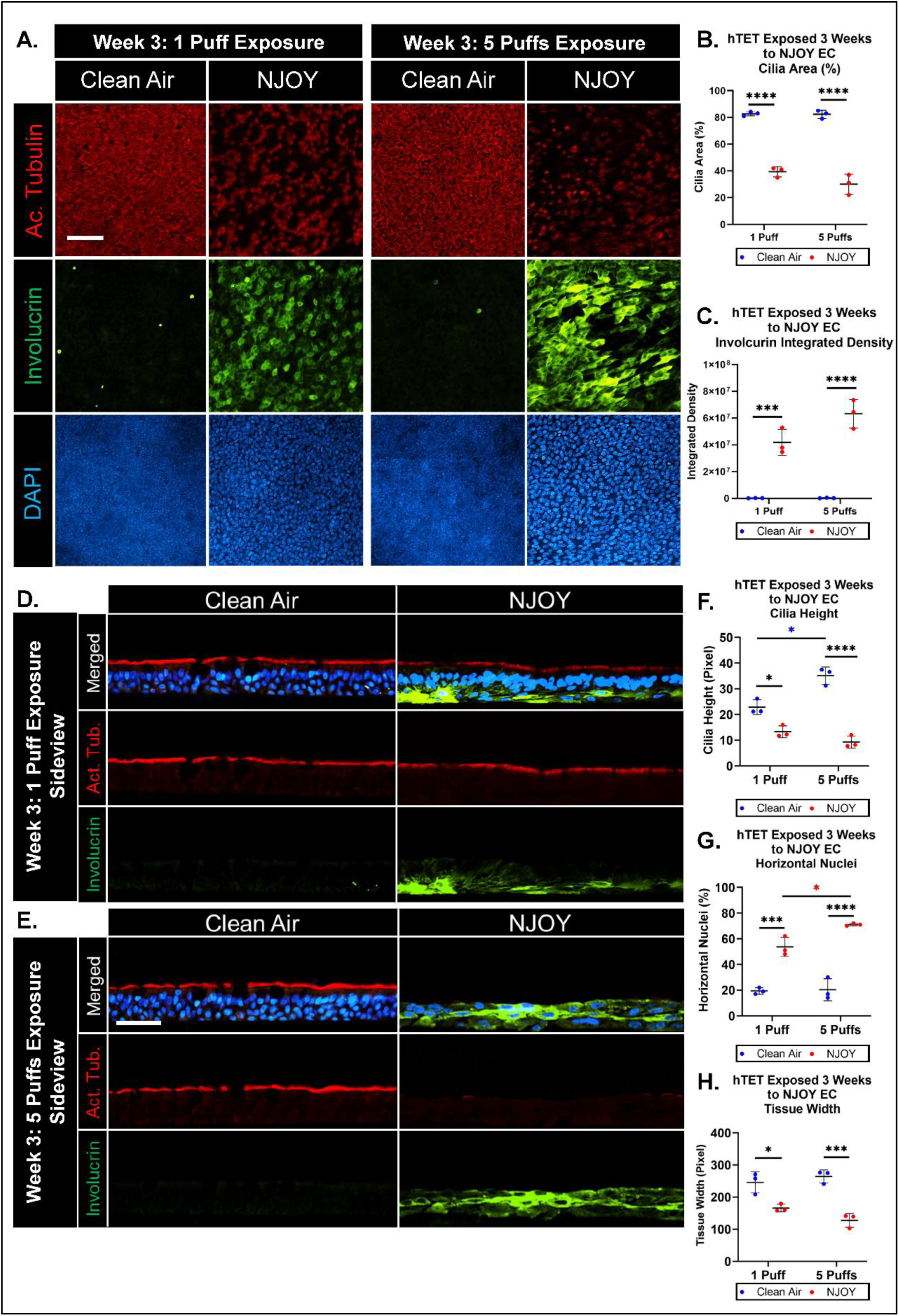
Three Weeks of NJOY EC Aerosol Exposure Induced Squamous Metaplasia in Normal Airway Epithelium. **(A)** BEV Immunofluorescence micrographs of hTETs exposed daily for 3 weeks to 1 (left) or 5 puffs (right) of clean air or NJOY EC aerosols. Tissues were stained with DAPI (nuclei; blue) and with antibodies to acetylated α-tubulin (cilia; red) or involucrin (green). Scale bar = 100 µm. **(B-C)** Quantification of **(B)** ciliary coverage and **(C)** involucrin fluorescence signal as integrated density. **(D-E)** Histological sections of hTETs exposed to 1 or 5 puffs of clean air or NJOY EC aerosol. Tissues were stained with DAPI (nuclei; blue) and with antibodies to acetylated α-tubulin (cilia; red) or involucrin (green). Scale bar = 50 µm. **(F-H)** Morphometric analysis showing **(F)** ciliary height, **(G)** percentage of horizontally oriented nuclei, and **(H)** tissue width. All graphs are the means ± standard deviations of three independent experiments. In **B, C, F, G,** and **H,** log(y) transformed data were analyzed using a one-way ANOVA followed by Tukey’s post hoc test to compare the means. Black * = significantly different than the control. Colored * significantly different between exposure groups (blue = clean air; red = NJOY). * = p < 0.05, ** = p < 0.01, *** = p < 0.001, **** = p < 0.0001.

Involucrin expression was markedly elevated in both EC aerosol exposure groups compared to clean air controls (Figure 4A), with involucrin localized to cells exhibiting squamous morphology. Quantitative analysis revealed that the integrated density of involucrin was significantly increased in both the 1- and 5-puff groups relative to their respective controls (Figure 4C). Furthermore, unlike week 2, involucrin levels were not significant in the 1- and 5-puff EC exposure group. Histological sections through hTET confirmed elevation of involucrin in both exposure groups (Figure 4 D, E).

In addition, the prevalence of horizontally oriented nuclei was significantly elevated in EC aerosol-exposed tissues compared to their respective controls, with a significant difference observed between the 1- and 5-puff groups (Figure 4G). Epithelial thickness was reduced in both aerosol exposure groups relative to controls, with the 5-puff group displaying greater, but not significant, thinning than the 1-puff group (Figure 4H).

Collectively, 3 weeks of NJOY EC aerosol exposure induced hallmark features of squamous metaplasia in hTETs, including ciliary loss, reduced ciliary height, increased involucrin expression, increased frequency of horizontally oriented nuclei, and diminished epithelial thickness.

### Comparison of Involucrin Levels in Weeks 1, 2, and 3

Involucrin levels were further compared after 1, 2, and 3 weeks of exposure using Western blot analysis on tissue lysates exposed to 1 or 5 puffs of clean air or NJOY aerosol (Figures 5A-5D). Clean air-exposed tissues in the 1- and 5-puff groups had similar involucrin levels, indicating a stable baseline.

**Figure 5:**
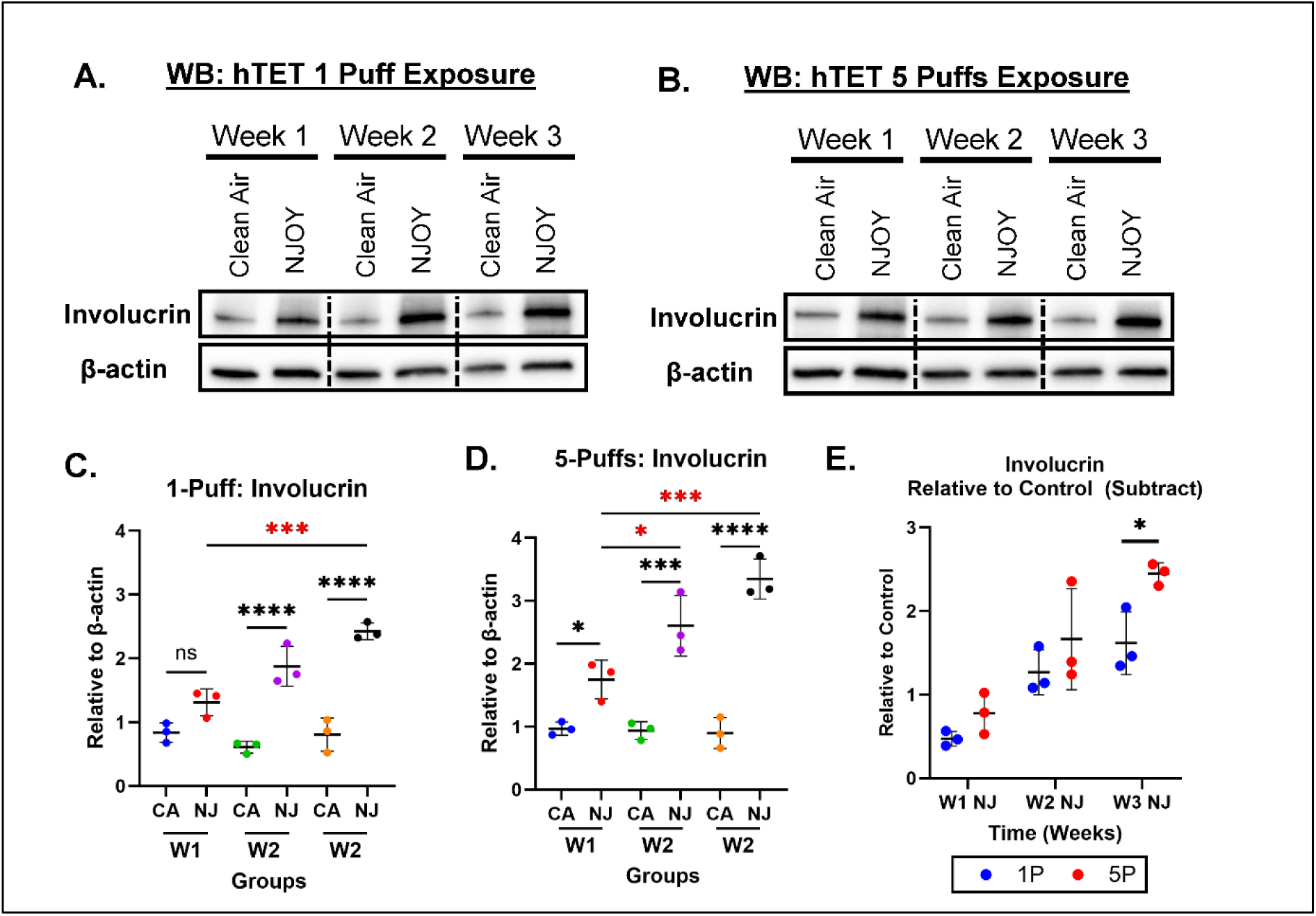
Western blot Analysis of Involucrin in hTET following EC Aerosol Exposure. **(A, C)** The effect of 1 puff of NJOY EC aerosol on involucrin levels in hTET following 1, 2, or 3 weeks of exposure. **(A)** A representative Western blot. **(C)** Involucrin levels relative to β-actin averaged from three Western blots. **(B, D)** The effect of 5 puffs of NJOY EC aerosol on involucrin levels in hTET following 1, 2, or 3 weeks of exposure. **(B)** A representative Western blot. **(D)** Involucrin levels relative to β-actin averaged from three Western blots. **(E)** Normalized involucrin levels (relative to clean air controls) at each and puff number. All graphs show the means ± standard deviations of three independent experiments. In **B, D** and **E, data** were analyzed using a one-way ANOVA followed by Tukey’s post hoc test to compare means. Black * = significantly different than the control. Red * = significantly different groups. * = p < 0.05, ** = p < 0.01, *** = p < 0.001.

At each progressive week, hTETs exposed to NJOY EC (1 or 5 puffs) aerosol showed significantly elevated involucrin concentrations compared to their respective clean air controls (Figure 5C and 5D). In both the 1- and 5-puff NJOY-exposed groups, involucrin concentrations were elevated relative to clean air controls and significant week-to-week differences were also observed, with involucrin levels rising progressively across the exposure period (Figure 5C and 5D).

To assess the effect of puff number on involucrin, the Western blot data were normalized to their respective clean air controls (Figure 5E). At each week, involucrin means were higher in the 5-puff group than the 1 puff group, and by week 3 this difference reached significance, supporting a concentration-dependent effect.

## Discussion

This study demonstrates that repeated ALI exposure of hTET to NJOY EC aerosol over 3 weeks induced progressive epithelial remodeling, beginning with goblet cell hyperplasia and culminating in SM. Remodeling was dependent on the number of puffs (1 vs 5) and the length of exposure (1, 2, or 3 weeks). Three weeks was sufficient for remodeling the hTET from a normal pseudostratified phenotype to an epithelium with the characteristics of SM. The main findings of the study are summarized in the graphical summary (Figure 6). The endpoints were quantifiable and amenable to statistical analysis. We propose that the hTET model presented here could become a standard assay used to study the effects of environmental chemicals, including EC aerosols, on the lining of the human respiratory system and provide data on a pathological change (SM) that may forecast more serious diseases, including COPD and squamous cell carcinoma. The data provided by this model could be valuable in estimating risk of specific exposures and in establishing regulatory policies for consumer products such as ECs. The hTET/ALI exposure model of SM provides numerous advantages over currently used assays. These include: the use of human tissues, the model is 3-dimensional, exposure is at the ALI, the endpoints are quantifiable, the experiment can be performed in a reasonable length of time, and the endpoints give a valuable readout on remodeling alterations that create a pathological state and forewarn of more serious consequences.

**Figure 6:**
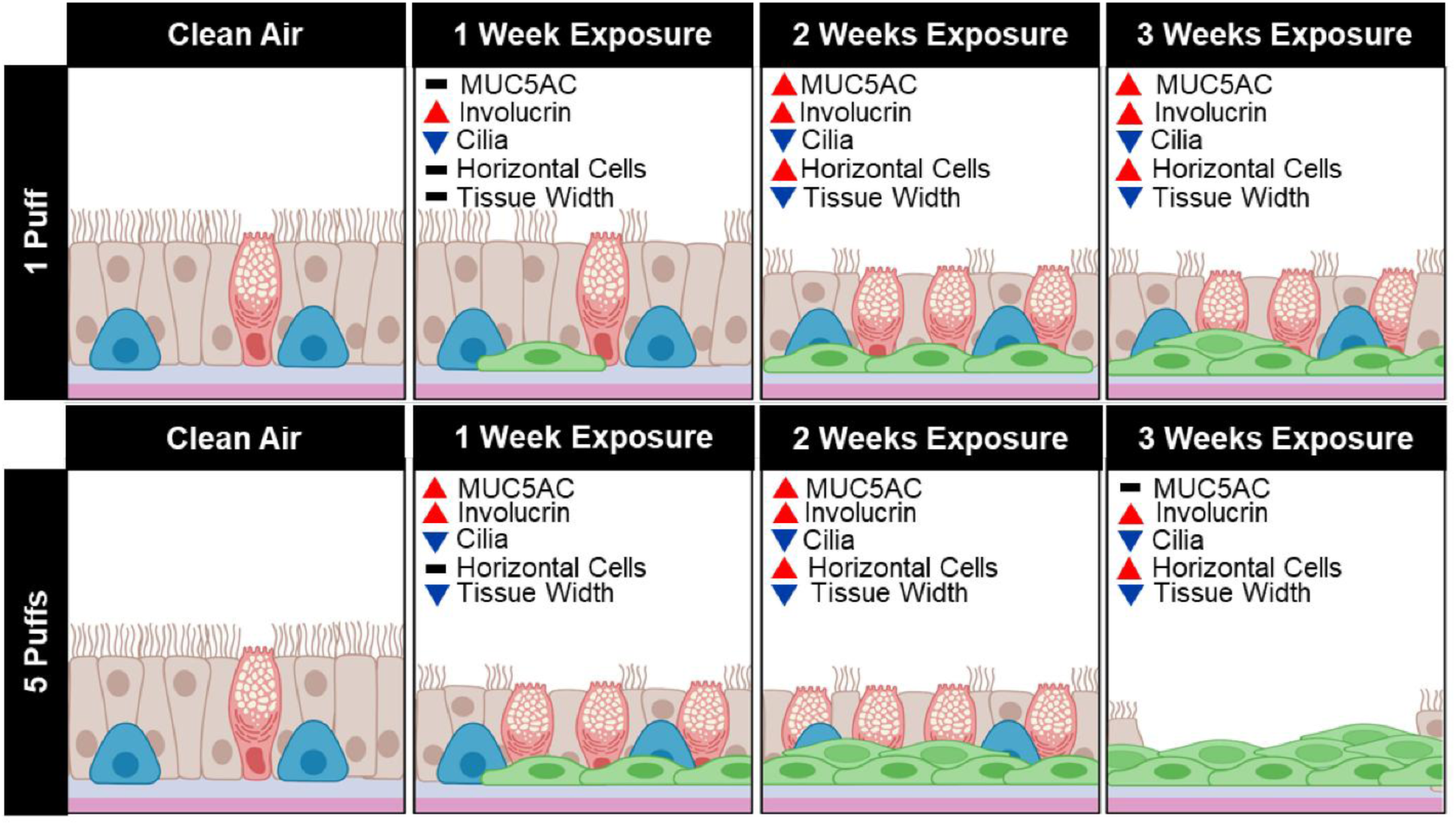
Graphical summary of EC aerosol induced epithelial remodeling in hTETs. Schematic summarizing the main findings of the study. Control tissues displayed a normal pseudostratified airway epithelium containing ciliated cells (brown), goblet cells (red), and basal stem cells (blue). Tissues were challenged with NJOY “Classic Tobacco” EC aerosol, exposed daily to either 1 or 5 puffs for 3 weeks, and analyzed at the end of weeks 1, 2, and 3. Quantitative endpoints include MUC5AC concentration, involucrin levels, ciliary coverage, percentage of horizontally oriented cells, and epithelial thickness. Black bars indicate no significant change compared with control, red upward arrows indicate a significant increase, and blue downward arrows indicate a significant decrease. The diagram depicts the progressive loss of mucociliary features and development of squamous characteristics (green cells) with increasing puff number and exposure duration.

To capture these progressive changes and model their relevance to human health, a 3-week exposure period was selected as it was long enough to capture remodeling, yet short enough to be experimentally practical. By generating both goblet cell hyperplasia and SM using controlled conditions, this system provides a reproducible and quantifiable framework for studying EC-induced airway remodeling and disease progression. It overcomes the limitations of animal models and short-term monolayer cultures and importantly further extends existing approaches by providing a direct quantitative read-out of human airway pathology.

Beyond capturing epithelial responses, the model provides a potentially powerful platform for risk prediction, IVIVE, and regulatory applications. By integrating multiple SM endpoints, it generates concentration and time-resolved data reflective of human airway biology. The endpoint data provide a mechanistic foundation for IVIVE, facilitating linkage between in vitro findings, clinical outcomes, and epidemiological trends. Such integration enhances the predictive capacity of experimental studies, enables early identification of airway injury markers, and informs the development of exposure thresholds aimed at mitigating long-term disease risk. Importantly, the model yields human-relevant risk data without relying on animal testing, therefore strengthening its translational utility for public health and regulatory decision-making.

This hTET platform also allowed us to characterize the sequence of epithelial responses over repeated exposures. Early responses to EC aerosol exposure included increased MUC5AC expression, consistent with a hypersecretory phenotype commonly observed in smokers and vapers [49] and reported in sputum samples from EC users [35]. In mouse models, EC aldehydes, such as acrolein, directly upregulated mucin expression [50]. The transient nature of this response, particularly the decline in MUC5AC levels by week 3 in the high-exposure group, demonstrates a shift away from goblet cell hyperplasia toward an alternative epithelial state. This transition was accompanied by significant reductions in ciliary coverage and height, increased involucrin expression, and elevated frequencies of horizontally oriented nuclei, all indicative of squamous differentiation.

Western blot analysis confirmed a cumulative increase in involucrin levels across exposure weeks, with a significant concentration-dependent elevation by week 3. Our findings align well with a prior transcriptomic study identifying SM markers in nasal biopsies from human EC users [38] and add both morphological and biochemical evidence that directly demonstrates concentration dependent progression toward SM in a human airway model exposed only to EC aerosols for a product with FDA authorization. Importantly, clean air exposures, even at the higher puff count, did not elicit comparable changes, underscoring the specificity of the EC aerosol effect.

Our model provides clinically relevant insights into the respiratory risks associated with EC use. By recapitulating key pathological features observed in EC users, specifically goblet cell hyperplasia followed by squamous metaplasia, this model provides quantifiable measures of airway injury that may precede overt clinical diagnosis. When considered with epidemiological studies showing increased risks of respiratory symptoms, COPD, and hypertension among EC users [3, 37], our findings help bridge experimental evidence with population-level outcomes. This translational connection reinforces the value of in vitro airway models in informing exposure thresholds, identifying high-risk usage patterns, and enabling earlier clinical intervention in respiratory care.

Our data show that EC aerosols activate pathways that contribute to squamous remodeling. San Juan et al. [51] reported that oxidative DNA damage can promote squamous transdifferentiation through p53 activation and G2/M checkpoint dysregulation. In line with this, EC aerosols induce reactive oxygen species accumulation, barrier dysfunction, and DNAdamage in bronchial epithelial cells [32], while also generating mutagenic DNA adducts and suppressing repair activity [52]. Taken together, these processes could provide a plausible pathway from exposure to the squamous features observed in our model, although further investigation is needed to directly establish these mechanistic links.

Given the public health significance of our findings, it is important to interpret them within the framework of current regulatory policies. NJOY ACE, the EC product evaluated in this study, is among the limited number of ECs granted premarket authorization by the U.S. Food and Drug Administration (FDA) under the Premarket Tobacco Product Application (PMTA) pathway [53]. This authorization, granted in 2022 for NJOY tobacco-flavored pods, was based on the determination that the product may confer a net public health benefit by offering adult smokers a potentially less harmful alternative to combustible cigarettes. However, FDA authorization does not constitute an endorsement of product safety, nor does it imply the absence of long-term or subacute health risks.

Our data demonstrate that even with relatively short-term exposures NJOY ACE aerosol induces hallmark features of SM, including ciliary loss, involucrin overexpression, and epithelial thinning. These findings underscore the need to expand regulatory toxicology frameworks to include histopathological and epithelial remodeling endpoints in 3D tissues, such as goblet cell hyperplasia, ciliary integrity, and SM, as part of premarket evaluations and post market surveillance. Incorporating these relevant pathological endpoints will strengthen risk assessment protocols and improve the detection of early airway injury associated with EC use.

## Conclusions

Our study identifies SM as a robust, measurable pathological outcome of EC aerosol exposure in human airway tissues. By providing time-resolved assessment, this model advances inhalation toxicology and offers a valuable quantitative platform for regulatory and preclinical safety evaluations of EC products. The induction of SM by EC aerosol exposure in a human airway model underscores the potential for vaping products to disrupt epithelial homeostasis and compromise respiratory defense mechanisms, such as goblet cell secretion. These findings carry significant translational weight, particularly in the context of rising EC use among adolescents and young adults. By establishing a mechanistically anchored, concentration-responsive model of epithelial remodeling, this study provides a valuable framework for evaluating the chronic toxicity of EC constituents and formulations. Moreover, the reproducibility and scalability of this system support its integration into regulatory toxicology pipelines, where it may inform product standards, risk assessments, and public health policy aimed at mitigating long-term respiratory harm.

## Abbreviations

EC: electronic cigarettes
hTETs: human tracheobronchial epithelial tissue
ABSC: airway basal stem cell
ALI: air-liquid-interface
SM: squamous metaplasia

## Acknowledgements

We thank The International Institute for the Advancement of Medicine (IIAM.org) for providing a human lung for isolation of the airway basal stem cells. We would like to thank the School of Medicine Research Core at UCR for providing some of the equipment that was used in this investigation. Figure 6 was created with BioRender.com.

## Authors’ contributions

Project administration and funding acquisition, P.T.; Conceptualization, R. P. and P.T.; Investigation R.P.; Sample preparation, data collection, and data processing were done by R.P., E.R., B.R., M.K., A.C., A.T. and T.M.; Data Interpretation was done by R.P. and P.T.; Writing the original draft was done by R.P. and P.T. Reviewing and editing was done by R.P., P.T. and T.M..

## Availability of data and materials

All data used in this study are contained in the manuscript.

## Ethics approval and consent to participate

Not applicable.

## Consent for publication

Not applicable.

